# TMS Over Dorsolateral Prefrontal Cortex Affects the Timing of Motor Imagery but not Overt Action: Further Support for the Motor-Cognitive Model

**DOI:** 10.1101/2022.02.25.481944

**Authors:** Marie Martel, Scott Glover

## Abstract

The Motor-Cognitive model suggests a functional dissociation between motor imagery and overt action, in contrast to the Functional Equivalence view of common processes between the two behaviours. According to the Motor-Cognitive model, motor imagery differs from overt action primarily through the use of executive resources to monitor and elaborate a motor image during execution, which can result in a lack of correspondence between motor imagery and its overt action counterpart. The present study examined the importance of executive resources in motor imagery by using TMS to impair the function of the dorsolateral prefrontal cortex while measuring the time to complete imagined versus overt actions. In two experiments, TMS over the dorsolateral prefrontal cortex slowed motor imagery but did not affect overt actions. TMS over the same region also interfered with performance of a mental calculation task, though it did not reliably affect less demanding cognitive tasks also thought to rely on executive functions. Taken together, these results were consistent with the Motor-Cognitive model but not with the idea of functional equivalence. The implications of these results for the theoretical understanding of motor imagery, and potential applications of the Motor-Cognitive model to the use of motor imagery in training and rehabilitation, are discussed.

## Introduction

Humans have a remarkable ability to mentally simulate their own actions. This ability, referred to as motor imagery, is used widely as a technique in motor training and skill learning (see for review [1,2]), as well as neurological rehabilitation (see for review [3–7]). Given its theoretical and applied value, it is important to develop and evaluate competing theories of motor imagery [8–10]. Two such models tested in the present study are the Motor-Cognitive model [11,12] and the Functional Equivalence view [13–15]

In the Motor-Cognitive model, both motor imagery and overt actions involve distinct planning and execution stages. Prior to initiation, the planning of both behaviours relies heavily on internal motor representations whereby an appropriate motor programme is selected based on information originating from the environment and the body and forms the basis of a forward model [16–20]. During execution, however, the processes underlying motor imagery and overt action diverge. In an overt action, predictions of the consequences of the movement are compared with sensory feedback obtained during its execution, and unconscious online sensory feedback is used to monitor and, if need be, correct the movement online [19,20]. In contrast, in the unfolding of motor imagery, no such sensory feedback is available. Instead, the execution of motor imagery is held to involve the conscious monitoring and elaboration of the unfolding motor image, a process that can depend heavily on executive resources [11,12].

A key prediction of the Motor-Cognitive model is that motor imagery should be highly sensitive to interference with executive resources, whereas overt actions should be relatively immune. This prediction has been upheld in previous work by Glover and colleagues [11,12]. There, participants were required to perform interference tasks heavily dependent on executive resources, including a calculation task involving counting backwards by threes, or generating words from a single root letter, while simultaneously executing or imagining an action. In line with the Motor-Cognitive model, both calculation and word generation lengthened imagined movement times much more than overt movement times [11,12]. Glover and colleagues argued that the interference tasks used executive resources normally engaged by the motor imagery system, resulting in the latter being slower in generating imagined movements.

Executive functions are heavily associated with the dorsolateral prefrontal cortex (DLPFC), which has been linked with such processes as directing attention, working memory, and inhibition [21–28]. If the Motor-Cognitive model is correct, DLPFC involvement should be important in motor imagery, and its involvement should be more important the greater the reliance on executive resources, such as in the motor imagery of actions for which detailed planning representations do not exist. This notion is consistent with previous studies. For one, numerous studies have observed a greater activation in the DLPFC during motor imagery than overt action [18,29–38], with this activation being even greater for novel movements [34,39]. For another, motor imagery training has been shown to improve working memory through an increase in prefrontal cortical activity [40], and coupling from Supplementary Motor Area to DLPFC is critical during motor imagery [41,42], with higher connectivity at resting-state predicting greater performance [43,44].

In contrast to the Motor Cognitive model, the Functional Equivalence view of motor imagery suggests a strong neural and behavioural correspondence between overt and imagined actions (e.g. [13,45,46]). Much early evidence supported this view. For example, both imagined and overt actions have been shown to follow Fitts law [35,45,47–50]. Both also responded similarly to biomechanical constraints imposed by a movement, such as adapting the body posture [51], correctly predicting inertial resistance [52,53], or adapting to the orientation of the object [54–56]. However, other studies observed numerous discrepancies between imagined and overt actions. For example, imagined movements can take either less time [57–60], or more time [11,12,48,57,60–66], than their overt counterparts. Further, macroscopic elements of imagined and overt movements may sometimes differ [67]. The predicted neural equivalence between motor imagery and overt actions in the Functional Equivalence model is also open to question (see for review, [31]). Although much overlap exists, numerous differences in the brain activity and connectivity of overt actions versus motor imagery are evident, including but not limited to the discrepancies in activity in the DLPFC noted above [41,68– 73].

In the present study we tested an important prediction of the Motor-Cognitive model that directly conflicts with the predictions of the Functional Equivalence view. Here, we used Transcranial Magnetic Stimulation (TMS) to disrupt the DLPFC in two experiments involving imagined and overt reaching and grasping tasks similar to those used by Glover and colleagues [11,12]. In the first experiment, participants had to execute and imagine executing a movement before and after receiving TMS or sham TMS. In the second experiment, participants had to either execute or imagine executing a movement after receiving TMS or Sham, with or without performing a simultaneous interference task designed to draw on executive resources.

In each experiment, we independently tested the effects of the TMS on the executive functions of working memory, inhibition, and task switching measured through a variety of computer-based cognitive tests. All these tasks have been noted to involve a large network of frontal areas in the brain [74–78], including the right DLPFC. In principle, as each of these tasks involves executive functions, if they are disrupted by TMS over the right DLPFC, then so too should motor imagery, assuming the latter also relies on executive functions as is argued by the Motor-Cognitive Model. More subtly, choosing a large range of tasks allowed us to investigate which, if any, specific executive functions might be most involved in motor imagery. For example, if TMS had deleterious effects on motor imagery, but only some of the cognitive tasks, this would suggest that the processes involved in the affected task(s) contribute heavily to the role of the DLPFC in motor imagery. We also tested the effects of TMS on performance in the calculation task of Experiment 2, used previously by Glover and colleagues [11,12]. Notably, calculation relies heavily on executive functions, as it involves a series of conscious mental operations, and activates the right DLPFC [74]. Thus, testing TMS effects on the calculation task would allow us to further investigate the type(s) of executive functions used in motor imagery.

Within the Motor-Cognitive framework, disrupting DLPFC activity should have deleterious effects on executive functions that slow the execution of motor imagery while leaving overt actions relatively unaffected. Conversely, within the Functional Equivalence framework, both overt actions and motor imagery ought to be similarly affected by all variables, including TMS over the DLPFC. The critical and novel test of the two theories thus revolved around the interaction between the variables of TMS and Action (overt action vs. motor imagery). To anticipate, in both experiments we observed that the application of TMS over the DLPFC affected motor imagery but not overt action, supporting the Motor-Cognitive model but not the Functional Equivalence view.

## Experiment 1

### Methods

#### Participants

We enrolled 32 participants (29 females, age range 18-32 years old) from the Royal Holloway Department of Psychology undergraduate research participation pool. All participants were right-handed by self-report, had normal or corrected-to-normal vision, had no motor or neurological impairments, and were naïve as to the exact purpose of the experiment. All passed the screening test for contraindications to TMS based on the guidelines from Rossi and collaborators [79]. Participants were randomly assigned to either the TMS or Sham group. Each participant gave written informed consent for the study, which was approved by the College Ethics Committee at Royal Holloway, and received course credits in return for participation. Three participants had to be replaced due to technical issues or because they failed to follow the instructions. Two further participants were excluded from the sample during data analysis (see Go/NoGo task section), leaving a total of 30 participants (28 females, age range 18-32 years old, 15 participants in each group).

Sample size was determined *a priori* and was similar to that in the two previous studies using the same tasks [11,12]. It was constrained by funding, which meant that the reliability of obtained point estimates (means) would be relatively low. However, given the very large effect sizes previously reported for the action tasks used here [11,12] we could be reasonably certain that these effects would replicate within even a limited sample.

#### Procedure

The procedure was organised into three main phases for a total duration of approximately 2h20 including consent form, setup, breaks and debrief (see Figure 1): First, a PRE-stimulation phase that involved completion of the cognitive and action tasks. Second, application of TMS/Sham over the DLPFC. Third, a repeat of the same cognitive and action tasks POST TMS/Sham. The cognitive tasks were always conducted in the following order: Spatial span, Go/NoGo and Switching. All participants took part in all phases of testing.

**Figure 1.**
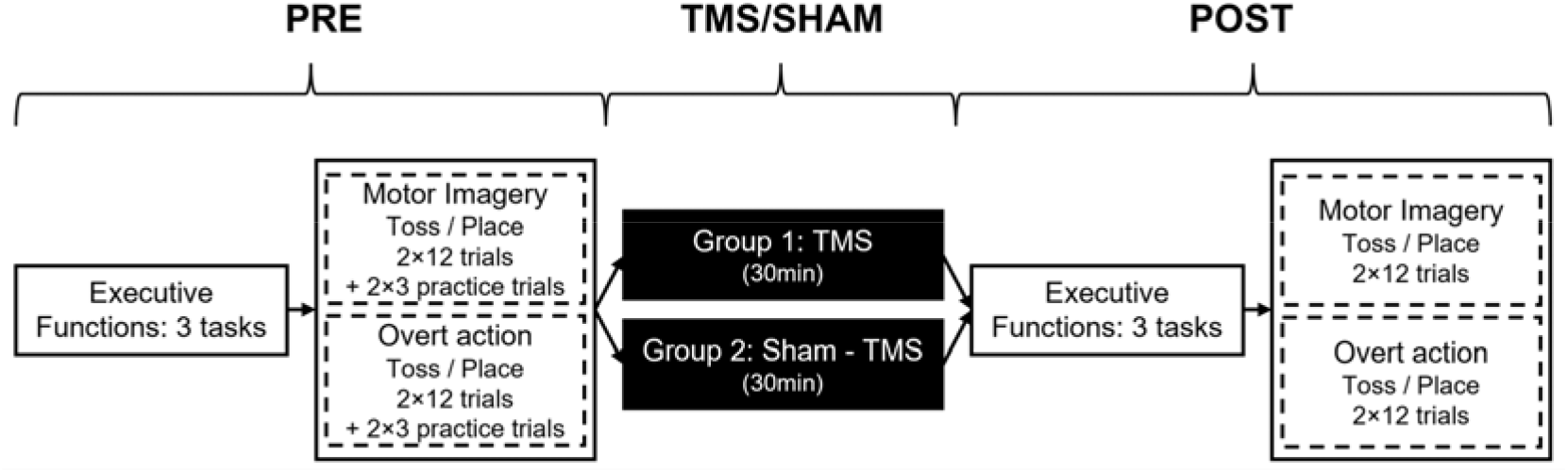
Experiment 1 Timeline.

Each participant performed both the overt action and motor imagery tasks, both PRE and POST TMS/sham, with order of task counterbalanced across participants following an ABBA design. For both action tasks, participants performed both “tossing” and “placing” actions (see below); the order of these was also counterbalanced using an AABB design.

#### Cognitive tasks

The three cognitive tasks included: A Corsi-style Spatial span task, a Go/Nogo task, and a Switching task. Instructions were displayed on the screen prior to each task (monitor size: 47.5 × 30 cm, resolution: 1680 × 1050 px). All tasks were implemented on PsychoPy3 (v3.1.5; [80]) adapted from the Psych Toolkit website (https://www.psytoolkit.org/). The full session of cognitive tasks took approximately 20-25 minutes.

##### Spatial span

The Spatial span task measured visuospatial working memory performance using a computerized version of the classic Corsi-block test [81]. The layout consisted of nine squares (size: 22 mm) displayed in an uneven pattern on the computer screen. At trial onset, individual squares were lightened at random (duration = 500ms, inter-stimuli interval = 250ms). Participants had to reproduce the correct sequence by clicking on the squares in the same order as just viewed with the mouse, and then clicking on a ‘Done’ button. Performance was not timed. The first sequence consisted of one square, with a square added to each ensuing trial following a sequence done correctly, and a square removed when participants failed to reproduce the sequence, up to a maximum of nine and minimum of one square. Different series of squares were used on each trial. Feedback was given on the computer screen for 2sec at the end of each trial. There were four practice trials followed by 16 analyzed trials.

##### Go/NoGo

The Go/NoGo task is designed to measure response inhibition [82]. It consisted of 12 practice trials, and 548 analyzed trials, with random stimulus presentations (476 Go and 84 NoGo stimuli, i.e. 85% and 15%, respectively,) with 600ms durations and random inter-trial intervals between 200ms and 600ms – these parameters followed the guidelines from Wessel [82] to increase the likelihood of prepotent motor responses. Participants had to press the space bar with their right index finger whenever a Go stimulus would appear (“GO – Press the space bar” written in black on a green oval) but refrain for NoGo stimulus (“NOGO – Press nothing” written in black on a red oval). A correct response for a Go trial was defined as a keypress within 600ms, whereas for a NoGo trial, it was defined by an absence of keypress within this time window. An incorrect response for a Go trial was an absence of a response within 600ms, whereas for a NoGo trial, an incorrect response occurred when the participant pressed a key within this time window. On incorrect trials, an error screen would appear for two seconds with a message defining the type of error (“You should not have pressed the button but you did” or “You should have pressed the button but you did not”). Feedback was added to ensure that participants would know when they made a mistake and to keep them focussed during the task.

##### Switching task

The Switching task measures cognitive flexibility [83]. It consisted of responding to one item in a combination comprised of one letter and one number based on the location of the stimuli. Each stimulus appeared in a 2 × 2 grid at the centre of the screen. If the stimulus appeared in one of the upper quadrants, participants were required to respond to the letter. If it appeared in one of the lower quadrants, they had to respond to the number. For responses to the letter, participants were required to press the “b” key with their left index finger for consonants, and the “n” key with their right index finger for vowels. For numbers, participants had to press the “b” key for odd numbers and the “n” key for even. There were 8 letters (A-E-I-U-G-K-M-R) and 8 numbers (2 to 9), for 64 possible stimulus combinations. Identical numbers or letters did not appear on consecutive trials.

The task was divided into three blocks, two “control” blocks of 64 trials each, followed by a “mixed” block of 128 trials. The first 12 trials of each control block and the first 12 trials of the mixed block were considered practice and not analysed; this left a total of 104 control trials and 116 mixed trials. In the first control block, the combinations appeared only in the upper quadrants and participants always responded to the letter. In the second control block, combinations appeared only in the lower quadrants and participants always responded to the number. In both blocks, lateral positions alternated in each trial between left and right.

In the final block, trials alternated between switching and no-switching trials. Here, stimulus position rotated through every position of the quadrant in a clockwise direction. The appropriate response thus switched on every second trial between letters and numbers (e.g., when the stimulus position moved from the top right to bottom right quadrant), but did not switch on the intermingled trials (e.g., when stimulus position moved from the bottom right to bottom left quadrant). In this block then, half of the trials were “mixed, no switch”, and half were “mixed, switch” trials. Participants had up to five seconds to respond. In case of an incorrect response, an error screen was displayed for three seconds with a reminder of the rules.

#### Action tasks

The experimental setup for the action tasks is shown in Figure 2. Participants sat comfortably at a 50 × 90 cm table set inside a large room. A computer keyboard, a wooden box (dimensions: 15.7 × 42 × 22 cm), and a small white plastic “poker chip” disc were on top of the table. The box contained a grey plastic tube (dimensions: 15 × 4.2 cm) positioned 18.5 cm from the participant’s end of the table, and the Polhemus receiver module, positioned in the corner of the box furthest from the participant. The starting position was a yellow square sticker (0.5 × 0.5 cm), positioned 52 cm away from the left side of the table, and 5 cm away from the near side. The disc was 1 cm thick and 4 cm in diameter. All other dimensions and positions were as shown in Figure 2.

**Figure 2.**
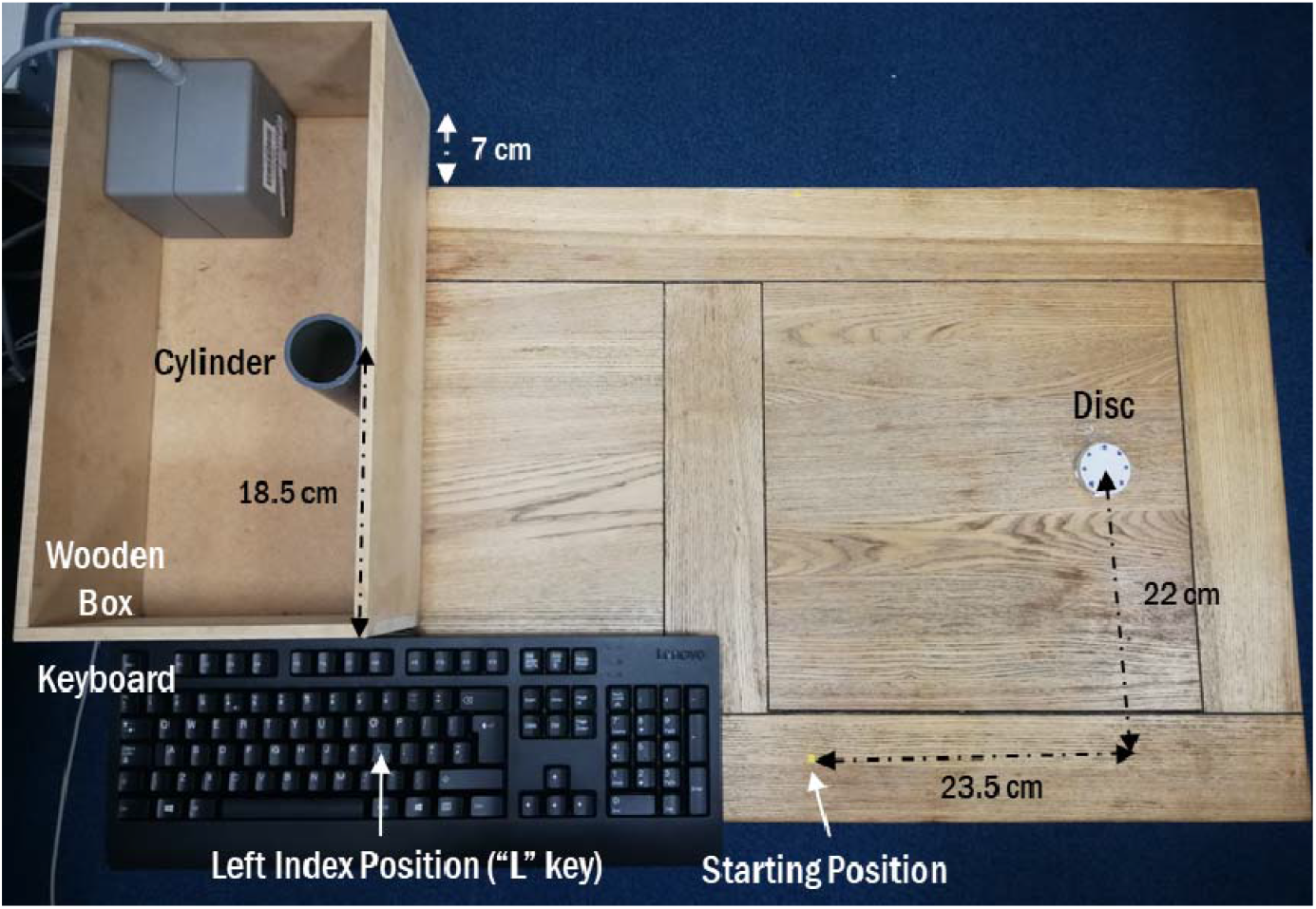
Experimental setup for the action tasks.

Before the start of the action tasks, two Polhemus transmitters (7 mm in diameter) were placed on the participant’s right thumb and index fingernails. Spatial location of these transmitters in x,y,z coordinates was recorded with a Polhemus Fastrak system (sampling rate: 120 Hz). For each movement, we extracted and analyzed the movement data off-line.

Participants sat with their body midline aligned with the starting position and began each trial with their right thumb and index finger held together in a comfortable “pinch” posture on the starting mark, and their left index finger resting on the “L” key. Each trial began with the experimenter triggering a tone via PC. In the overt action block, participants had to either reach for the disc and place it into the cylinder (“place” – high precision) or reach for the disc and toss it into the wooden box (“toss” – low precision). For the motor imagery block, participants had to imagine the same actions as vividly as possible, using both visual and kinaesthetic imagery of their movement experienced in the first person (i.e., to imagine seeing and feeling themselves performing the movement), with their eyes open, while keeping their right arm still. In both overt action and motor imagery sessions, participants pressed the “L” key at the beginning (or imagined beginning) of their movement, and again when releasing (or imagining releasing) the disc into the target area. During the overt action trials, the experimenter monitored participants to ensure they pressed the key at the appropriate times. In between trials, the experimenter put the disc back in place if needed and participants returned to the starting position (or remained in the same position in the motor imagery trials). Once the participant was set, the experimenter triggered the tone to begin the next trial. The action tasks session took approximately 15 minutes.

#### TMS

Prior to applying TMS (or Sham-TMS), we determined motor threshold for each participant, defined as the minimum intensity of TMS required to result in visible twitches of the finger following stimulation of the index finger/thumb area of the right primary motor cortex. Testing stimulus intensity was set at 90% of the individual’s motor threshold. Stimulation site for the right DLPFC was determined using the Beam method [84]. In short, we entered the distance tragus to tragus, nasion to inion, as well as the head circumference, for each participant. The software then indicated the position of the DLPFC (corresponding to the F3 electrode given by the 10-20 system) to be measured from the midline and the vertex. For the TMS group, TMS was applied over the stimulation site at 1 Hz for 30 min, for a total of 1800 pulses. For the Sham group, sham TMS was administrated to the same area by inverting the coil so that the magnetic field projected away from the scalp instead of through it. Due to the length of testing, the coil tended to overheat roughly twice per session, leading to a total TMS/Sham session of approximately 40min. Mean stimulation intensity was very similar between the two groups (TMS: 58.6%, sham: 57.8% of stimulator output, respectively).

#### Data processing

##### Action tasks keypress data

The first 3 trials of each task in the PRE session were considered practice. For the remaining trials, we extracted reaction time (RT) and movement time (MT) as recorded by the keypress data for both overt action and motor imagery. RT was the time between the starting tone and the first press of the key (indicating the beginning of the movement/imagery). MT was the time between first and second keypress. Any trial with RT or MT outside ± 2 IQR (Interquartile Range) for each combination of task, session and participant was removed from the analysis (3.8% of the total trials).

##### Kinematic data

Data obtained via the Polhemus was used to perform a manipulation check that participants were pressing the “L” key at times closely corresponding to their actual movement onset and offset. For the kinematic data, the beginning of the movement was set as the point at which a minimum speed of 0.05 m/s was obtained, and three different cut-off criteria were used for the end of the movement based on velocity or grip aperture. We then compared the reaction times and movement times using these criteria to those as indexed by the keypresses in the overt action task. Reaction and movement times calculated using these cut-offs were all highly correlated with the times based on keypresses (all r^2^ ≥ 0.70, see Supplementary Information).

##### Spatial span

For the sixteen experimental trials, we extracted the mean span score, computed using the formula from an adapted version of the Corsi paradigm [85]. This formula took into account both correct and incorrect responses to provide a representative score.

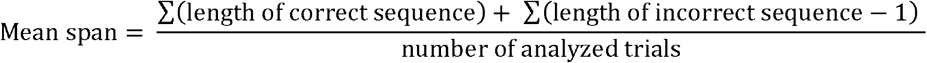

##### Go/NoGo

For the 548 experimental trials, we computed the mean error rates. First, trials with RT ≤ 200 ms were removed as anticipation errors (2.3% of total trials). Any participant with more than 10% anticipation errors in her overall performance was excluded from the experiment (2 participants). For the remaining trials, we computed the proportion of correct trials separately for Go and NoGo trials. For the analysis, the arcsine-transformed proportions were used as these provide a means of meeting the assumptions of normality, heterogeneity, and non-additivity required for parametric tests [86].

##### Switching

For the 220 experimental trials, we calculated the proportion of correct responses, then subjected these to an arcsine transformation for analysis. We then calculated the reaction time defined as the time between stimulus presentation and the participant’s response. Trials without a keypress were removed (0.1% of the trials). RTs outside ± 2 IQR (Interquartile Range) for each combination of task, session and participant were removed from the analysis (3.6% of the remaining trials).

#### Statistics

All raw data and analyses from both experiments are available here: https://osf.io/s82v4/. Analyses were performed with R Studio (R version 3.6.1; [87]).

The action tasks used a 2 × 2 × 2 × 2 × 2 mixed design with *TMS* (TMS vs. SHAM) and *Order* (OA first vs. MI first) as between-subjects variables, and *Session* (PRE vs. POST), *Action* (OA vs. MI), and *Precision* (Place vs. Toss) as within-subjects variables. We present analyses of the movement times based on keypresses. As explained in the Introduction, the critical tests revolved around the variables *TMS* and *Action*. Effects involving *Precision* were also expected to be present based on previous work [11]. The effects of the variables *Order* and *Session* and their interactions with other variables were not always relevant to distinguishing between the two theories under examination here, and as such, we treated these variables as exploratory. As we did not have *a priori* hypothesis for the reaction times, and there was no clear theoretical interpretation we could apply to any such effects, our exploratory analyses of RTs are presented separately in the Supplementary Information.

The Spatial span and the Go/NoGo tasks used a 2 × 2 mixed design with *TMS* (TMS vs. SHAM) as a between-subjects variable and *Session* (PRE vs. POST) as a within-subjects variable (for the Go/NoGo task, these analyses were run separately on the Go and NoGo trials). The Switching task used a 2 × 2 × 3 mixed design with *TMS* as a between-subjects variable and *Session* and *Trial Type* as within-subjects variables. *Trial Type* had 3 levels: “control” trials in which the same rule was used on every trial (letter only and number only blocks), “mixed – no switch” trials in which the rule was the same as the previous trial, and “mixed – switch” trials in which the rule had switched from the previous trial.

Following Glover and Dixon [88], we report the adjusted likelihood ratios (λ_adj_) [89] of nested linear models calculated from the sums-of-squares tables obtained via ANOVA. The use of likelihood ratios is a philosophical approach to statistical inference based on comparing the relative strength of competing statistical models, each of which can be directly linked to a theoretical account: A likelihood ratio is the relative likelihood of the data given two possible statistical models. Likelihood ratios avoid many of the well-known pitfalls associated with *p*-values and NHST [90–93], and unlike Bayesian statistics, make no prior assumptions about the probability distribution of different hypotheses but report the evidence based on the data alone. Critically, the use of likelihood ratios in combination with nested linear models allows for directly testing complex models that include multiple effects and interactions to simpler models, tests which cannot be done straightforwardly using *p*-values. In contrast, *p*-values, which represent the probability of the observed effect or something larger occurring under one statistical model (the null), do not speak directly to the likelihood of the data given two competing hypotheses, but instead seek to control long-term error rates of false-acceptance or false-rejection of the Null. As a more complex model will nearly always provide a better fit to the data, we used adjusted likelihood ratios that penalise models with a greater number of parameters based on the Akaike Information Criterion [89] adjustment. In many prototypical hypothesis-testing situations, likelihood ratios bear a close relationship with *p*-values, and λ_adj_ = 3 will generally correspond to *p* ∼ 0.05, whereas λ_adj_ = 10 will generally correspond to *p* ∼ 0.02.

The approach of comparing nested models means that effect(s), interaction(s), and/or specific contrasts (referred to collectively as “variables” in the remainder of this paragraph) are added to statistical models, either singly or in groups, prior to each statistical test. The choice of which variables to add will be largely motivated by the theories under consideration. For the data from the action tasks for example, we first tested models which included variables consistent with both the Motor-Cognitive and Functional Equivalence view; this was mainly to illustrate the presence and strength of the evidence for these effects. We then went on to the critical tests of the two theories that included effects of variables predicted only by one or the other theory; this allowed us to determine which theory the data was more likely to occur under. We completed these analyses by comparing the fits of a statistical model including all of the variables that were solely predicted by the Motor-Cognitive model to one that included only those variables consistent with Functional Equivalence. Some exploratory tests were conducted as well that included variables not of direct theoretical interest. Tests of data from the cognitive tasks were generally more straightforward and were conducted mainly to assess the likelihood of TMS having affected performance, as that was the main interest in the analysis of those tasks.

### Results

#### Keypress movement times

We first examined effects consistent with both the Motor-Cognitive and Functional Equivalence views. A statistical model including effects of *Session, Precision*, and the *Session × Precision* interaction fit the data much better than did a null model (λ_adj_ > 1000). These results showed that movement times as indexed by the keypresses were longer in the high-precision condition (placing) as compared to the low-precision condition (tossing), were shorter in the POST session than in the PRE session, and that the decrease in MT from PRE to POST sessions was greater for placing than for tossing (Figure 3A). All these effects were consistent with both the Motor-Cognitive and Functional Equivalence views.

**Figure 3.**
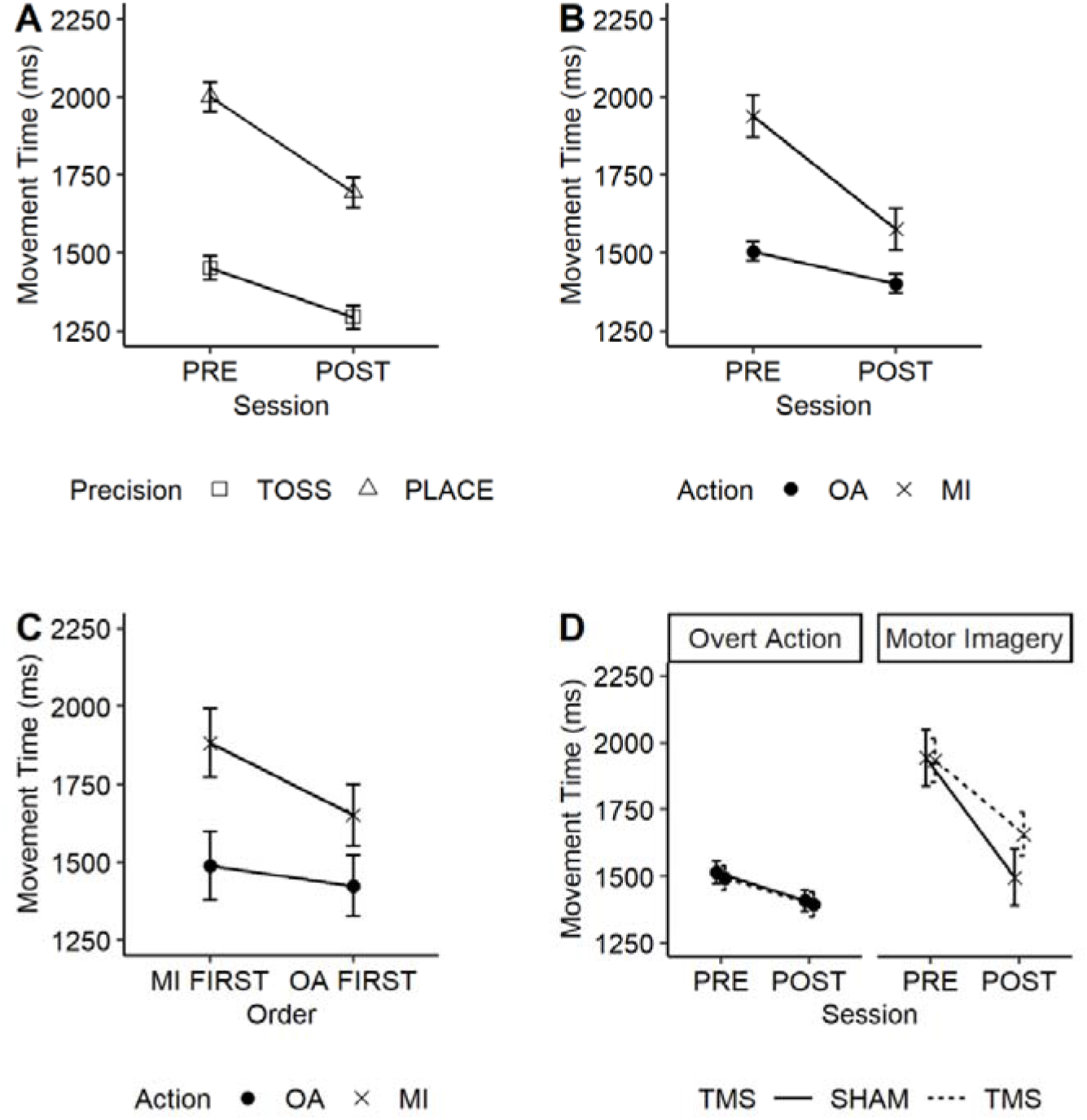
Depiction of interactions of interest tested in the different models. Panel A: Effects of Session, Precision, and Session × Precision on movement times. Panel B: Effect of Action in the PRE and POST sessions. Panel C: Effect of Order on overt action and motor imagery keypress movement times. Panel D: Effects of TMS on overt action and motor imagery keypress movement times in Sham and TMS groups in the PRE and POST sessions. In panels A, B and D, error bars represent the standard errors of the pre-post difference and are appropriate for within-subject comparisons. In panel C, error bars represent the standard errors of the MI-OA difference and are also appropriate for within-subject comparisons. Refer to Figure S1 in the supplement for the overall effects of the variables of *Session, Precision, Action*, and *TMS* on keypress movement times in the action tasks, with individual data points.

We next explored effects predicted uniquely by the Motor-Cognitive model. Adding a main effect of *Action* whereby movement times were longer in motor imagery than in overt action improved the fit immensely (λ_adj_ = 552.6). Adding a *Session* × *Action* interaction further improved the fit (λ_adj_ = 90.4), indicating that movement times were overall more similar between motor imagery and overt action in the POST session than in the PRE session (Figure 3B). The fit was further improved greatly by adding a complex contrast in which the effect of *Action* was modulated by *Order*, but only for motor imagery, relative to a model that predicted *Order* effects that were independent of *Action* (λ_adj_ > 1000). This contrast showed that movement times for motor imagery were shorter when overt action was performed first, whereas performance in overt action was unaffected by having done motor imagery first (Figure 3C). All these effects were consistent only with the Motor-Cognitive model and not the Functional Equivalence view.

Most critically, the fit was further improved by adding a contrast whereby the effects of TMS were limited to motor imagery trials in the POST session (λ_adj_ = 7.10; Figure 3D). This was as predicted by the Motor-Cognitive model. However, the evidence did not support the fit of a contrast whereby the effect of *Precision* was greater in the motor imagery than in overt action (λ_adj_ = 1.16), as also predicted by the Motor-Cognitive model and had been observed in a previous study [11].

Finally, to summarize the relative ability of the Motor-Cognitive and Functional Equivalence models to predict the keypress MT data, we compared a statistical model including all the factors predicted by the Motor-Cognitive model (*Session, Precision, Session* × *Precision, Action, Session* × *Action*, and contrasts involving interactions between *Action* × *Order, Action* × *TMS*, and *Action* × *Precision*) to one including only the factors consistent with both theoretical views (*Session, Precision, Session* × *Precision*). The former model provided an immensely better fit (λ_adj_ > 1000). In other words, the data as a whole were over a thousand times more likely to occur when including those effects uniquely predicted by the Motor-Cognitive model than when including only those effects consistent with both the Motor-Cognitive and Functional Equivalence models.

#### Cognitive Tasks

##### Spatial span

A model assuming an effect of *Session* (PRE vs. POST) fit the data much better than did a null model (λ_adj_ = 268), showing that performance overall improved with practice. Adding an interaction *Session* × *TMS* improved the fit fairly modestly (λ_adj_ = 2.2), suggesting that the TMS may have led to slightly less improvement following practice relative to Sham stimulation (mean ± SD; Sham: 5.1 ± 0.6 vs. 5.7 ± 0.5 in POST; TMS: 5.0 ± 0.7 vs. 5.3 ± 0.7 items in POST). This suggested, albeit tentatively, that working memory may be a critical component of executive functions in motor imagery.

##### Go/NoGo

Mean accuracy for the different conditions in the Go/NoGo task are shown in Table 1. For Go trials, the evidence somewhat favoured the null over a model including effects of *TMS, Session*, and *TMS × Session* (λ_adj_ = 6.51). For the NoGo trials, the best-fitting model included only an effect of *Session* in which accuracy decreased with practice (PRE vs. POST; λ_adj_ > 1000). In sum, there was no evidence that TMS negatively affected performance in the Go/NoGo task. Thus, it seemed unlikely that inhibition plays a major role in motor imagery.

**Table 1.** Accuracy in Go and NoGo trials before and after the (Sham) TMS session.

|  | Mean accuracy in Go trials (%) |  | Mean accuracy in NoGo trials (%) |  |
| --- | --- | --- | --- | --- |
|  | PRE | POST | PRE | POST |
| Sham | 94.7 | 95.1 | 77.1 | 68.4 |
| TMS | 95.6 | 95.6 | 74.6 | 68.8 |

##### Switching – Accuracy

Data from the switching task are shown in Table 2. The best-fitting model for the proportion of correct responses included only an effect of *Trial Type* (λ_adj_ > 1000), indicating reduced accuracy for the Mixed-Switch trials. Adding the main effect of *TMS* and its interaction with *Trial Type* slightly worsened the fit (λ_adj_ = 0.54, or λ_adj_ = 1.87 in favour of the simpler model). In short, there was no evidence that TMS affected accuracy in this task.

**Table 2.** Accuracy and reaction times in the task-switching.

| | Mean accuracy (%) | | | | Mean reaction times ± SD (ms) | | | |
| --- | --- | --- | --- | --- | --- | --- | --- | --- |
|  | Sham |  | TMS |  | Sham |  | TMS |  |
|  | PRE | POST | PRE | POST | PRE | POST | PRE | POST |
| Control | 96.3 | 93.7 | 95.4 | 95.5 | 649 ± 118 | 574 ± 56 | 631 ± 63 | 590 ± 46 |
| Mixed – No Switch | 95.6 | 95.2 | 96.6 | 96.4 | 880 ± 236 | 722 ± 165 | 843 ± 168 | 727 ± 111 |
| Mixed – Switch | 92.5 | 93.4 | 91.3 | 92.3 | 1175 ± 398 | 933 ± 297 | 1177 ± 299 | 929 ± 195 |

##### Switching – Reaction Times

The best-fitting model included effects of *Trial Type, Session*, and the *Trial Type* × *Session* interaction (λ_adj_ > 1000), in which participants were overall faster in the control task, faster in the POST session, and the reduction in RT across sessions was greater in the more difficult trials (Mixed – No switch and Mixed – Switch). This model fit better than a model that also included effects of TMS and its interaction with session (λ_adj_ = 5.76), again suggesting TMS did not affect performance. From this, it appeared that cognitive switching did not underly the use of executive functions in motor imagery.

### Discussion

Keypress movement time data from the action task strongly supported the Motor-Cognitive-model over the Functional Equivalence model. Critically, TMS was associated with longer movement times in the motor imagery task only, leaving overt actions unaffected. Practice between PRE and POST improved the correspondence between the timing of motor imagery and overt action to a greater extent in the Sham Group than the TMS group, reinforcing the idea that disrupting the DLPFC disrupted the ability to perform motor imagery effectively. Of some interest, our exploratory tests of order effects found that whether overt action was performed before or after motor imagery affected movement times. This may have arisen because with physical practice a better representation of the movement becomes available to motor imagery [65].

Results of the cognitive tasks were inconsistent. TMS may have slightly disrupted performance in the spatial span task but did not appear to affect performance in the Go/NoGo or switching tasks. This could suggest many possibilities: Most parsimoniously, it may be that working memory is the executive function most important to motor imagery, as suggested by a study showing improvement in working memory after motor imagery training [40], and that inhibition and/or task switching are less important, or not important at all. More speculatively, it may be that motor imagery shares with all these tasks a common pool of executive resources, and that the cognitive tasks did not use enough of these executive resources to be clearly affected by TMS. We address these possibilities in greater detail in the General Discussion.

In Experiment 2, we simplified the design and increased the sample size to focus on a smaller number of key variables, remove exploratory variables that might conceivably confound the results, and gain more precise estimates of effects. Relative to Experiment 1, we eliminated the variables *Order, Session*, and *Precision*, and added the variable *Calc* [for calculation]. The lattermost variable was introduced as it has previously been shown to have a large impact on the timing of motor imagery with only a minimal effect on overt action [11,12]. It also represents a classic example of a task that relies heavily on executive resources [74]. The removal of the exploratory variables of *Order, Session*, and *Precision* in Experiment 2 that were not of direct theoretical interest strengthened our design as this eliminated the possibility of these variables introducing spurious effects or spurious interactions that might mask main effects. The pre-registered methods, hypotheses, and analysis plan of Experiment 2 can be viewed here: https://osf.io/geqx2.

Once again, the two theories make contrasting predictions. According to the Motor-Cognitive model, only motor imagery ought to be affected by the *Action, TMS* and *Calc* variables, and by any interactions between *Action* and either or both of the other two variables. Further, the slowing of movement times by TMS should be greater when the calculation task is performed concurrently with motor imagery, evident in a three-way *Action × TMS × Calc* interaction. Conversely, the Functional Equivalence model predicts none of the above: no main effect of *Action*, nor any interactions between *Action* and either of the *TMS* or *Calc* variables, nor a three-way *TMS* × *Action* × *Calc* interaction.

## Experiment 2

### Methods

#### Participants

We enrolled 64 new participants (58 females, age range 18-55 years old) from the Royal Holloway Department of Psychology undergraduate research participation pool and research staff. Note that there was only one participant aged 55 years, with in total two participants older than 32, which makes our two experiments comparable in terms of age. Two participants were replaced due to technical difficulties. One participant was further excluded from the sample during data analysis (due to high anticipation error rate in the Go/NoGo as in Experiment 1), leaving a total of 63 participants (57 females, age range 18-55 years old). Participants were randomly assigned to receive either TMS (n=31) or Sham-TMS (n=32). Participants in each of these groups were further randomly assigned to take part in either the motor imagery (n = 16 in each group) or the overt action condition (TMS: n = 15; Sham-TMS: n = 16). Inclusion criteria were the same as in Experiment 1. Due to Covid-19 safety precautions, the last 25 participants performed all tasks with a face mask and surgical gloves on. None of the participants reported any additional difficulty in executing the tasks with these safety measures. These latter participants received a £15 retail voucher instead of credits. The sample size was again based on funding constraints, but was increased relative to Experiment 1 to provide more reliable point estimates.

#### Procedure

The entire experiment lasted about 1h30 per participant, including consent form, setup, breaks and debrief. Participants first received TMS (or sham TMS) at a rate of 1Hz for 30 minutes (1800 pulses) over the right dorsolateral prefrontal cortex. Immediately after TMS/Sham stimulation, they comfortably sat in front of a computer to perform the three cognitive tasks, following the same order as in Experiment 1: Spatial span, Go/NoGo, and Switching (about 20-25 min). Following this, participants performed either motor imagery or overt action of the grasping and placing task from Experiment 1 (about 15 min). Each participant performed two blocks: one of the two blocks for each participant included the calculation task, the other was a control without the calculation, counterbalanced across participants using an ABBA design (see Figure 4).

**Figure 4.**
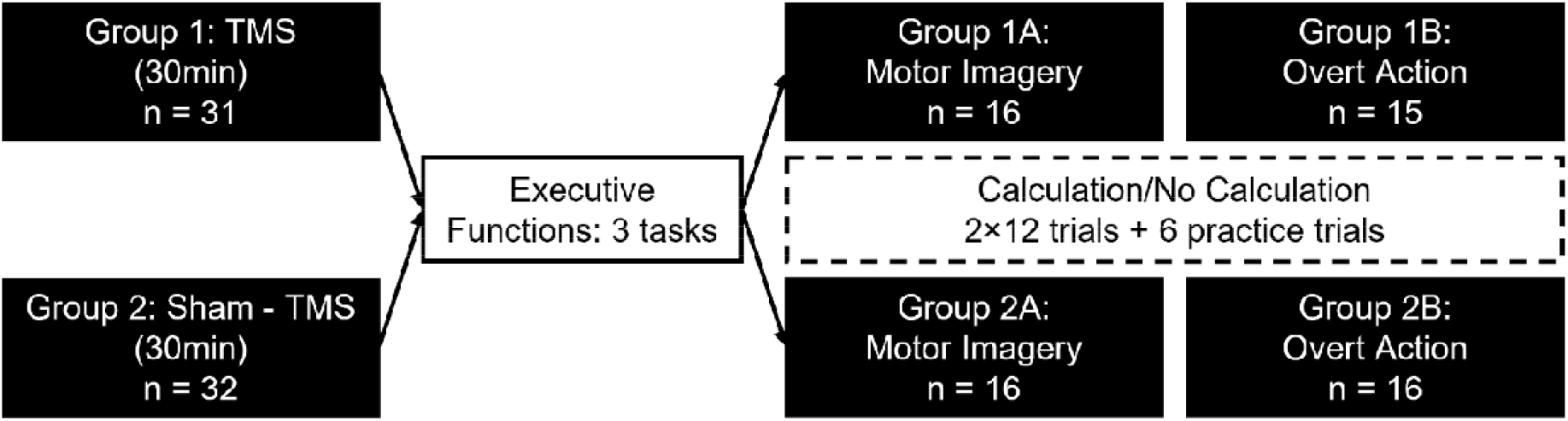
Experiment 2 timeline. Participants first received TMS or Sham TMS, then performed the three executive function tasks, then performed either the motor imagery or overt action version of the action task, each of which included calculation and no-calculation trials.

#### Cognitive tasks

The cognitive tasks were identical to Experiment 1.

#### Action tasks

The action task setup was identical to Experiment 1. As in Experiment 1, participants indicated the beginning and end of each movement (or its imagined counterpart) by pressing the “L” key on the keyboard. For the motor imagery group, participants had to imagine the action as vividly as possible, using both visual and kinaesthetic imagery of their movement experienced in the first person (i.e., to imagine seeing and feeling themselves performing the movement), with their eyes open, while keeping their right arm still. In the calculation task, just prior to the onset of the trial, the experimenter read aloud a pre-determined number between 51-99. At the sound of the tone, participants had to simultaneously count backwards out loud by threes while performing either the overt or imagined reaching and placing movement. The numbers were pseudo-randomly determined so that the same multiples of three could not occur in successive trials (as might occur if, e.g., the numbers 63 and 60 were given on two adjacent trials). Verbal responses were recorded by the experimenter for later analysis.

#### TMS

The application of TMS/Sham was identical to Experiment 1. Mean stimulation intensity was very similar between the TMS and Sham groups (TMS: 57.0%, Sham: 56.8% of stimulator intensity, respectively).

#### Data processing and statistics

Extracted variables, outlier detection and general data processing was as in Experiment 1, and were described in the preregistration of Experiment 2 (https://osf.io/geqx2). The action tasks were analysed using a 2 × 2 × 2 mixed design with *TMS* (TMS/Sham) and *Action* (overt action/motor imagery) as between-subjects variables, and *Calc* (calculation/control) as a within-subjects variable. As a control, we also conducted an analysis including a fourth variable, *TimelineCovid*, which separated participants based on whether or not they had to follow safety measures such as wearing a mask and gloves. As this analysis provided strong evidence that the *TimelineCovid* variable did not modulate any of the other effects (see Supplementary Information), we excluded this variable from further analyses.

Additionally, we analysed the performance in the Calculation task. The number of errors (i.e., wrong numbers) on the Calculation task was computed and entered into a 2 × 2 mixed design with *Group* (TMS/Sham) and *Action* (overt action/motor imagery) as between-subjects variables. We had originally planned on investigating the average number of responses for the calculation task (in the pre-registration) but later realized it would not be informative as a longer trial would allow more responses than a shorter trial, so instead we computed a “time per response”. This was indexed as the total number of responses given by a participant for each condition divided by the sum of movement times for each condition.

The Spatial span and the Go/NoGo tasks had a between-subjects design with *TMS* (TMS or SHAM) as the between-subjects variable, and the Switching task added the within-subjects variable *Trial Type* (control, mixed-no switch, and mixed-switch). As before, we report the fit of nested linear models using adjusted likelihood ratios, λ_adj_.

### Results

#### Keypress movement times

As in Experiment 1, we focussed our tests on the competing predictions of the Motor-Cognitive and Functional Equivalence models. Although the tests described here fulfilled this goal, note that the present analysis diverted somewhat from the pre-registered plan which had been based on predicted effect sizes, as the formulation of said plan turned out to include miscalculations which we only noticed after we had collected the data. In lieu of the pre-registered analyses, the analyses undertaken were the same conventional and statistically sound tests of nested linear models used in Experiment 1.

We again began by testing statistical models consistent with both theoretical views. A model including a main effect of *TMS* whereby movement times were longer in the TMS group than in the Sham group fit the data somewhat better than a null model (λ_adj_ = 3.66). There was also strong evidence for both a main effect of *Calc* and a *TMS × Calc* interaction wherein movement times were longer in the calculation condition, particularly for the TMS group (λ_adj_ > 1000). These effects strongly suggested that TMS interfered with subsequent performance of the calculation task, and were consistent with both the Motor-Cognitive and Functional Equivalence views (Figure 5A).

**Figure 5.**
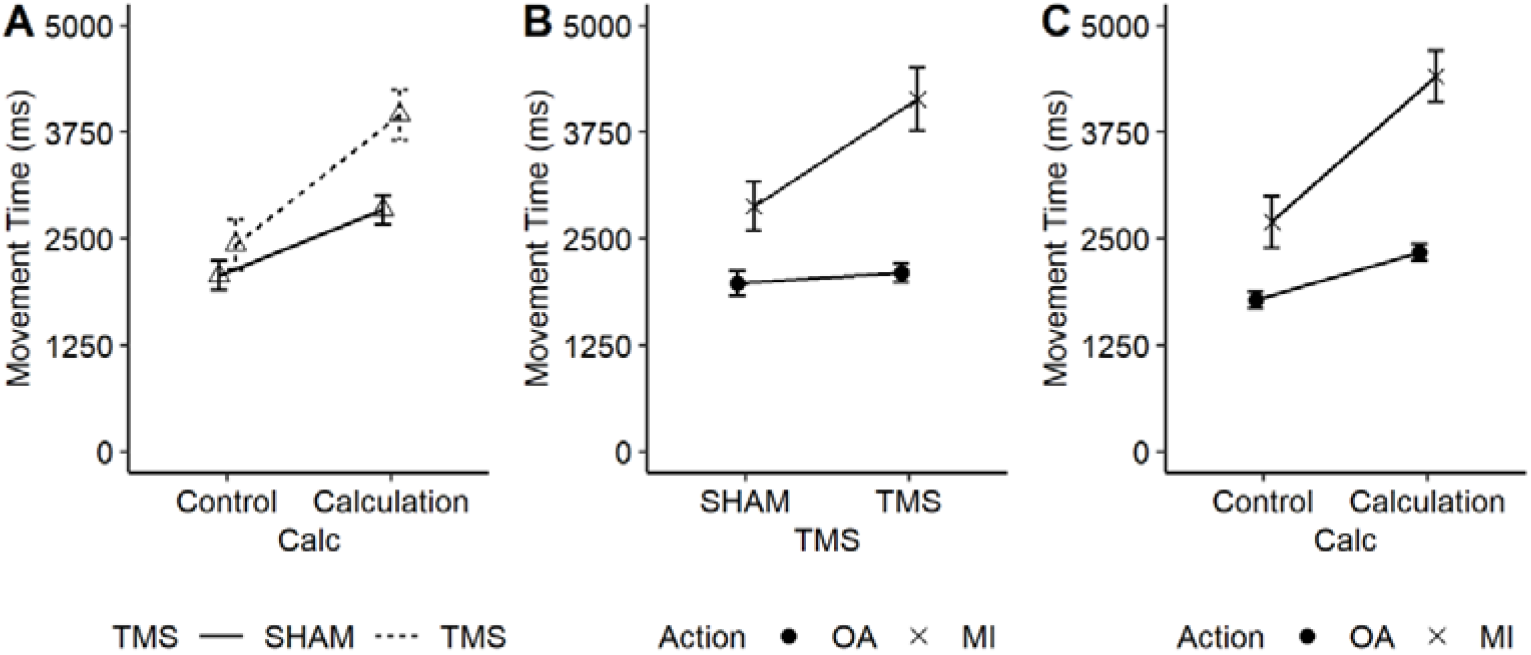
Depiction of interactions of interest in Experiment 2. Panel A: Effect of TMS on movement times under the control (no calculation) condition or with the calculation task. Panel B: Effect of Action in the TMS and Sham-TMS groups. Panel C: Effect of Action on movement times as a function of calculation task. In panels A and C, error bars represent the standard errors of the Calculation-Control difference and are appropriate for within-subject comparisons. In panel B, error bars represent the standard errors of the subject means and are appropriate for between-subject comparisons. Please refer to Figure S4 in the supplement for the full effects of the *Action, Calc*, and *Group* variables on keypress movement times in the overt and imagined reaching and grasping task.

We next examined effects predicted solely by the Motor-Cognitive model. Adding the main effect of *Action* and the *Action* × *TMS* interaction, improved the fit greatly (λ_adj_ > 1000). This provided strong evidence that movement times were longer overall in the motor imagery group than in the overt action group, and longer still in the TMS group in the motor imagery condition compared to the overt action condition (Figure 5B). Adding in the *Action* × *Calc* interaction, wherein movement times were longer in the calculation condition than in the control condition, and with this effect being greater in the motor imagery condition, improved the fit even more (λ_adj_ = 226; Figure 5C). However, there was little evidence to support adding the three-way interaction *Action* × *Calc* × *TMS*, also predicted solely by the Motor-Cognitive Model; including this interaction did not notably affect the fit (λ_adj_ = 1.03).

Finally, we again compared a statistical model including all the effects uniquely predicted by the Motor-Cognitive model (*Action, Action* × *TMS, Action* × *Calc*, and *Action* × *Calc* × *TMS*) to one including only those effects consistent with both Motor-Cognitive and Functional Equivalence views (*TMS, Calc, TMS* × *Calc*). This analysis showed strong support for including those effects unique to the Motor-Cognitive view (λ_adj_ > 1000). In short, the data were more than 1000 times as likely assuming the effects predicted solely by the Motor-Cognitive model than assuming only those effects predicted by the Functional Equivalence view.

#### Calculation Task

Time-per-response and error rates for the calculation task are summarised in Table 3. For the mean time–per-response, the best-fitting model included main effects of both *TMS* and *Action* (λ_adj_ = 102.1). This indicated that participants took more time to calculate a number after TMS stimulation compared to Sham, and during the motor imagery compared to overt action condition, confirming that both TMS and motor imagery negatively affected performance of the calculation task. However, for error rates a null model fit the data better than a full model incorporating effects of *TMS, Action*, and *TMS* × *Action* (λ_adj_ = 9.2), suggesting none of these variables affected accuracy. The effects of *TMS* and *Action* on time per response, together with the lack of similar effects on error rates, indicated that the former did not result from participants employing a speed-accuracy trade-off.

**Table 3.** Calculation task performance.

| | Mean Time Per Response ± SD (ms) | | Error rate (%) | |
| --- | --- | --- | --- | --- |
|  | MI | OA | MI | OA |
| Sham | 1780 ± 530 | 1486 ± 658 | 4.56 | 7.56 |
| TMS | 2741 ± 1343 | 1831 ± 993 | 6.98 | 7.54 |

#### Cognitive Tasks

##### Spatial span

Spatial span score in Experiment 2 was explained slightly better by a null model that assumed no difference between TMS (5.3 ± 0.8 items) and Sham (5.2 ± 0.7 items) than by a model including an effect of *TMS* (λ_adj_ = 2.42). Thus, unlike in Experiment 1, the evidence modestly suggested TMS did not affect performance of this task, and that instead working memory may not be a critical executive function used in motor imagery.

##### Go/NoGo

Accuracy performance here was also similar between TMS and Sham groups in the Go trials, with a null model assuming no difference between TMS (95.1**%)** and Sham (95.5%) again providing a slightly better account of the data than a model assuming an effect of *TMS* (λ_adj_ = 2.38). For the NoGo trials, the evidence did not distinguish between the null and a model including the effect of TMS (λ_adj_ = 1.08 in favour of the null; TMS: 77.3%; Sham: 73.6%). These results confirmed those of Experiment 1 in suggesting that motor imagery did not heavily depend on inhibition.

##### Switching – Accuracy

Table 4 shows mean accuracy and reaction times for the switching task. The best fitting-model included a large effect of *Trial Type* (λ_adj_ > 1000) whereby accuracy was lower in the mixed-switch trials than in the control or mixed-no switch trials. However, adding a main effect of *TMS* and an interaction between *TMS* and *Trial Type* did not improve the fit at all (λ_adj_ = 1.0), suggesting that TMS did not affect accuracy.

**Table 4.** Accuracy and reaction times in the task-switching paradigm.

| | Mean accuracy (%) | | Mean reaction times ± SD (ms) | |
| --- | --- | --- | --- | --- |
|  | Sham | TMS | Sham | TMS |
| Control | 95.3 | 96.2 | 661 ± 86 | 683 ± 88 |
| Mixed – No switch | 97.6 | 96.6 | 811 ± 149 | 917 ± 188 |
| Mixed – Switch | 93.6 | 92.7 | 1152 ± 261 | 1223 ± 292 |

##### Switching – Reaction Times

The best fitting-model included a large effect of *Trial Type* (λ_adj_ > 1000) whereby reaction times were higher the more difficult the task was. Adding in a contrast whereby TMS slowed reaction times in the mixed-no switch and mixed-switch trials, but not in the control trials, did not notably improve the fit (λ_adj_ = 1.26). Again, these results were consistent with those of Experiment 1 in suggesting that set-switching was not an important executive function used during motor imagery.

### Discussion

The results of Experiment 2 provided further support for the Motor-Cognitive model over the Functional Equivalence view. Critically, as in Experiment 1, TMS had much greater effects on keypress movement times for motor imagery than for overt action. This suggested that disrupting

DLPFC activity with TMS interfered with the timing of motor imagery, but left overt actions relatively unaffected. Similarly, in line with previous studies [11,12], the calculation task also had a slowing effect on motor imagery but not on overt action. Overall, a statistical model that incorporated all the effects predicted solely by the Motor-Cognitive model performed much better than one including only those effects predicted by both the Motor-Cognitive and Functional Equivalence views.

TMS also had large effects on performance of the calculation task. Specifically, there was strong evidence that TMS lengthened the amount of time required to generate each number in the calculation task, relative to sham stimulation. Evidence for effects of TMS on the other three cognitive tasks, however, was inconclusive. Unlike in Experiment 1, TMS did not appear to affect performance of the spatial span task, and for the Go/NoGo and switching tasks, there was again no evidence that TMS impacted their performance. We discuss the possible reasons for these results in detail in the General Discussion.

### General Discussion

#### Support for the Motor-Cognitive Model

The results of the two experiments here strongly support the idea, inherent in the Motor-Cognitive model, that the DLPFC plays an important role in motor imagery. When DLPFC activity was disrupted by TMS, keypress movement times became longer in motor imagery tasks relative to sham stimulation, but the same effect was not observed in the overt action tasks. These findings are consistent with several studies using fMRI showing increased DLPFC activation during motor imagery [18,29–38]. However, they contradict a core principle of the Functional Equivalence view, which holds that motor imagery utilizes the same internal processes as overt actions, and thus both motor imagery and overt action should be equally affected (or unaffected) by disruption of the DLPFC.

The present study also replicated many of the main findings of Glover and colleagues [11,12]. First, motor imagery of the reaching, grasping and placing/tossing actions consistently took longer than the corresponding overt actions, independent of TMS or other manipulations. This we attribute to the extra cognitive load of indexing the timing of the imagined movements by pressing the keyboard, which utilises executive resources [11,12]. Second, adding an interference task greatly lengthened the time required to complete the motor imagery task, but had much smaller effects on overt actions. We attribute this to the use of executive resources in the calculation task, depriving motor imagery of these same resources. Replication of these findings provide further support to the Motor-Cognitive model over the principle of Functional Equivalence.

##### TMS and Cognitive Tasks

The effects of TMS on the calculation task and the three other cognitive tasks were inconsistent. These are important results to try to disentangle, as the logic of the design was that TMS over DLPFC would disrupt executive functions, which in principle ought to have affected all tasks thought to use these functions. Although TMS had a very large and unequivocal effect on the time required to give responses in the calculation task of Experiment 2, its effects on the other cognitive tasks were not generally evident; only weak evidence for an effect of TMS on the Spatial span task of Experiment 1 was observed, and this was not replicated for the same task in Experiment 2.

There are several possible explanations for why this pattern of results may have arisen. Each of the cognitive tasks we employed taps into different types of executive functions, and in fact one reason why we employed a wide range of tasks was to explore which of these functions might ally most with the functions used to support motor imagery. For example, spatial span performance is largely dependent on spatial working memory, which could presumably also be important in motor imagery. However, working memory is a largely distributed process that relies not just on the right DLPFC [76,77], but a large network of areas, including the left DLFPC [21,74], the ventral prefrontal cortex [94], the precentral sulcus [95] and the posterior parietal lobes [96–98]. Thus, disrupting the right DLPFC alone may have been insufficient to reliably affect spatial working memory. Performance in the Go/NoGo task depends on response inhibition, a process that has been argued to play an important role in motor imagery [99,100]. On this account, motor imagery must inhibit the pre-planned motor response and instead elaborate it mentally. Although we agree that there must be some element of inhibition in motor imagery, we did not find effects of TMS on Go/NoGo performance. It may be that the inhibitory element of the Go/NoGo task does not depend on the right DLPFC; rather it may rely on other prefrontal regions such as the left DLPFC, or more ventral regions of the prefrontal cortex [75,78].

Switching task performance has been attributed to at least two not mutually exclusive processes: working memory to select and maintain tasks sets, and a process to reduce interference from stimulus ambiguity [101,102]. These functions, activating frontal areas in the brain [103] require cognitive flexibility and executive resources, and thus could arguably be matched with processes supporting motor imagery, and so expected to be impaired by TMS over the right DLPFC. Yet here, no such effects were observed.

Given the generally absent effects of TMS on these three cognitive tasks, it may seem surprising that it did have large effects on the calculation task. As an internally-generated response, the calculation task may require a greater amount of executive resources than the other three cognitive tasks [104–106]. It also appears to be introspectively more demanding of mental resources than the other tasks. Perhaps not coincidentally, motor imagery is also internally-generated and is also introspectively a fairly demanding mental task. Further to this, performing arithmetic heavily activates the right DLPFC [74]. Each of these factors may have contributed to the TMS affecting performance of motor imagery and the calculation task, but not the other three cognitive tasks. In short then, the three other cognitive tasks we used to measure the effects of TMS over the DLPFC on executive functions may have been focussed fairly specifically on individual executive functions such as working memory, inhibition, and task switching, and less reliant on the general pool of executive resources. Cognitive tasks requiring internally-generated responses might indeed be more sensitive to TMS over the DLPFC, tasks such as the calculation task used here, word generation [12] generating novel uses of a common object, or reciting the months of the calendar in reverse order.

##### Practical Applications of the Motor-Cognitive Model

Given the support that has been found for the Motor Cognitive model here and in other recent work [11,12], it is worth exploring its potential applications in training and rehabilitation. Current techniques rely heavily on the notion of functional equivalence [107,108]. But if the Motor Cognitive model is a nearer approximation of the organisation of motor imagery, this could have important implications for the treatment of many motor and cognitive disorders, as well as the typical or atypical development of motor skills. At present, and while acknowledging the need for further testing of the model in the context of training and rehabilitation, our sense is that applying the Motor-Cognitive model to these pursuits will involve trainers and therapists considering the important role of executive functions in motor imagery, and the conditions which modulate their use.

In training and rehabilitation, for example, the Motor-Cognitive model would suggest that the best outcomes for motor imagery training would be those for which a strong motor representation of the action is already present. This would result in a higher fidelity between the initial formulation of the motor image and its corresponding overt action during the planning stage, resulting in a lower reliance of motor imagery on inefficient (in terms of resources) and low fidelity (in terms of macroscopic and microscopic movement elements) executive functions to elaborate the motor image [11]. Consistent with this view, completely novel movements for which no internal motor representation exists cannot be learned with motor imagery alone [6,109]. For novel or otherwise poorly represented actions, a better approach might be to use action observation to help build internal representations [110,111], which can then be strengthened with motor imagery.

A common motor disorder that may respond positively to incorporating the Motor-Cognitive model into its treatment is Developmental Coordination Disorder (DCD), in which persons experience impaired motor skills from an early age, negatively impacting academic and life achievements [112]. Consistent with the Motor-Cognitive model, these individuals also present with reduced motor imagery abilities and deficits in executive functions [113,114]. Treatment of DCD may potentially benefit from incorporating the tenets of the Motor-Cognitive model as suggested above – i.e., the first step towards improvement of motor function ought to be the building and strengthening of internal motor representations, either through overt practice or action observation, which may then be strengthened further with motor imagery training.

Patients with damage to the DLPFC also report poor motor imagery abilities [115]. A prediction of the Motor-Cognitive model is that these deficits should be most evident for imagined actions which rely most heavily on executive resources, dependent on the DLPFC. If this can be confirmed empirically, then rehabilitation efforts for patients with DLPFC involvement might follow the same guidance as above, perhaps even more strenuously, as those with DLPFC damage should be less able to mentally elaborate a motor image for which only a poor internal representation exists.

## Conclusion

The present study demonstrated an important role of the dorsolateral prefrontal cortex in motor imagery. When TMS was used to disrupt the right DLPFC, participants took longer to perform motor imagery compared to a sham-TMS condition. TMS had a similar effect on a calculation task designed to utilise a general pool of executive resources. TMS did not, however, reliably affect cognitive tasks more focussed on specific executive functions. These results support the idea, inherent in the Motor-Cognitive model, that the DLPFC plays an important role in motor imagery, and provide further evidence against the notion of a functional equivalence between motor imagery and overt actions. This may have important implications for the use of motor imagery in training and rehabilitation.

## Acknowledgments

This work was supported by the BIAL foundation (Grant 193/18). We thank Jai Patel, Emily Durber, Mollie Barker and Vykintas Maziukas for their help with data acquisition.

## Data availability statement

All data, code and material used in this study are available on the website of the Open Science Framework (OSF) at the following link: https://osf.io/s82v4/

## CRediT Author Statement

**Marie Martel:** Conceptualization, Methodology, Software, Formal analysis, Investigation, Data Curation, Writing - Original Draft, Writing - Review & Editing, Visualization, Supervision. **Scott Glover:** Conceptualization, Methodology, Software, Formal analysis, Resources, Writing - Review & Editing, Supervision, Project administration, Funding acquisition.

