## Supplementary Information for "TMS Over Dorsolateral Prefrontal Cortex Affects the Timing of Motor Imagery but not Overt Action: Further Support for the Motor-Cognitive Model"

1. Experiment 1
   1.1. Analysis of keypress movement times – Full figure

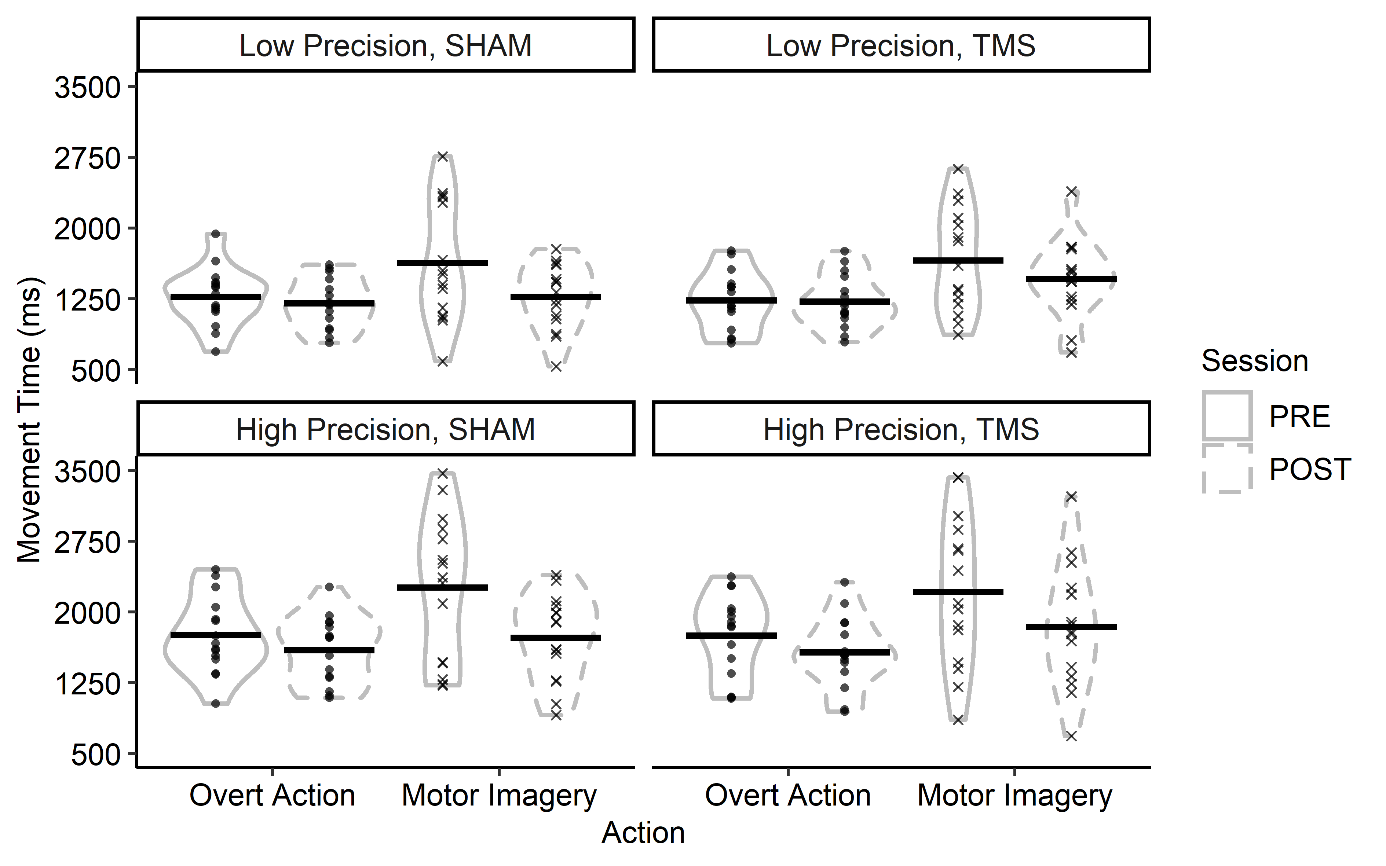

**Figure S1.** Effect of TMS on overt action (circle) and motor imagery (cross) movement times, for the low precision (tossing) and high precision (placing) tasks in Sham and TMS groups in the PRE (solid) and POST (dashed) sessions. Individual points represent the mean score for single participants and horizontal lines represent the group means. Enclosures represent the distribution of the presented data, with wider horizontal sections indicating a larger number of observations.

1.2. Kinematic data

In the main manuscript, we used keypresses as an indicator of the beginning/end of the movement in both MI and OA. Yet this approach is reliable only if keypresses are truly a reflection of the movement. We thus assessed if this was the case by investigating the correlation between keypress data and kinematics.

*Data processing.* The first 3 trials of each task in the PRE session were considered practice and thus removed from the analysis. We extracted the RT and three MT based on different kinematic criteria and compared them to the RT and MT data from the keypresses in the overt action condition. RT was calculated based on a standard velocity criterion whereby movement onset was defined as a minimum speed of 0.05 m/s. Movement times were computed using three different cut-offs: 1) minimum velocity; 2) constant increase of the maximum grip aperture; and 3) maximum grip aperture (MGA), all in the placing/tossing phase of the movement. Similarly to the keypress data, any overt action trial with RT or MT outside ± 2 IQR (Interquartile Range) for each separate task, session and participant was removed from the analysis (4.1% of the total trials). As the Matlab program extracting these values was automatized, closer investigation of the data was done if large variability was noted between button-pressing and kinematic measures. We then applied some cut-offs and discarded any trials outside the range (4.9% of total trials): 0<MT>5000ms; MT based on MGA criteria – MT based on velocity > 1500ms; MTs based on MGA criteria differed from more than 1000ms; MGA occurred at 150% of the MT or more. This ensured that the values extracted by the program were not artefacts.

*Statistics.* We calculated Pearson correlations for each participant and each MT criterion.

*Results.* There were generally large positive correlations between the keypressing indicating the start of the movement and the start of the movement based on the kinematic measure (RT), as well as between the keypressing indicating the end of the movement and the time of the end of the movement given by our three different kinematics markers (MTvel, MTinc and MTmga; Figure S1). Inspection of individual trials indicated that most of the participants pressed the key just before or after movement onset and again, just before or after movement completion.

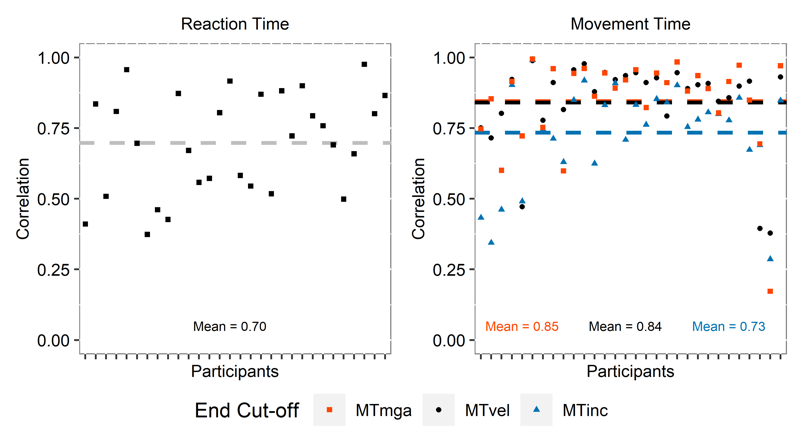

**Figure S2.** Pearson correlation values between the keypress data and the kinematics parameters in Experiment 1. Symbols are individual correlation values for the reaction times (left panel) or the movement times (right panel). In the right panel, MT values on the same column belong to the same participant. Dashed lines indicate the mean correlation for each marker. MT = Movement Time, defined by 3 different criteria: lowest velocity (vel), increase grip aperture (inc) or maximum grip aperture (mga) during the second part of the movement.

1.3. Analysis of keypress reaction times

Neither the Motor-Cognitive model nor the Functional Equivalence view make any specific predictions regarding the RT. We thus conducted an exploratory analysis on the data, these are presented in Figure S2. A model including an effect of *Session* whereby RT was lower in the POST than in the PRE session fit the data better than a null model (λ_adj_ = 47.5). Adding an effect of *Precision* wherein RT was longer in the high precision (placing) task than in the low precision (tossing) task had a very moderate effect (λ_adj_ = 2.3), while adding the *Session* × *Precision* interaction did not improve the fit (λ_adj_ = 0.48, or λ_adj_ = 2.07 in favour of a model without the interaction).

A model including a main effect of *Action* immensely improved the fit (λ_adj_ > 1000), which was further improved by including the interaction *Action* × *Session* (λ_adj_ = 87.8). This showed that participants had greater RTs for motor imagery compared to overt action, and that RT became smaller in the POST session but mostly for motor imagery, while RT for overt action was unaffected.

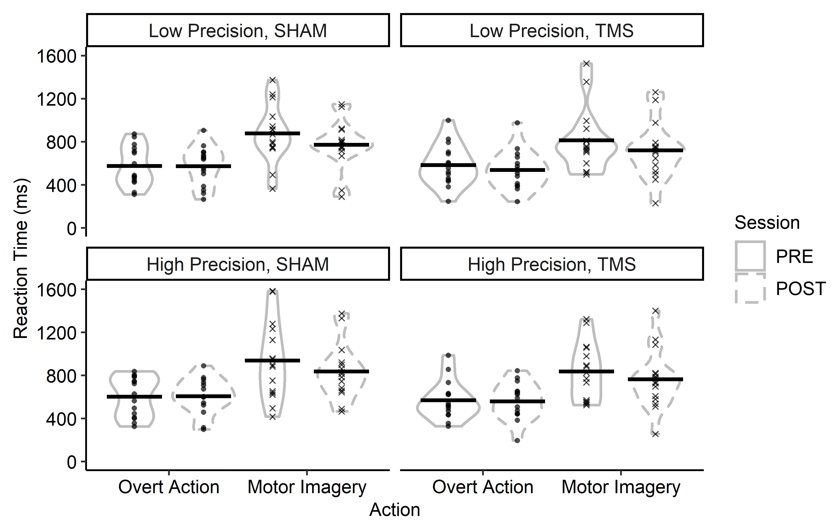

**Figure S3**. Effect of TMS on reaction times for overt action (circle) and motor imagery (cross), for the low (tossing) and high (placing) precision tasks in Sham and TMS groups in the PRE (solid) and POST (dashed) sessions, Experiment 1. Individual points represent the mean score for single participants and horizontal lines represent the group means. Enclosures represent the distribution of the presented data, with thicker sections indicating a larger number of observations.

2. Experiment 2
   2.1. Analysis of keypress movement times – Full figure

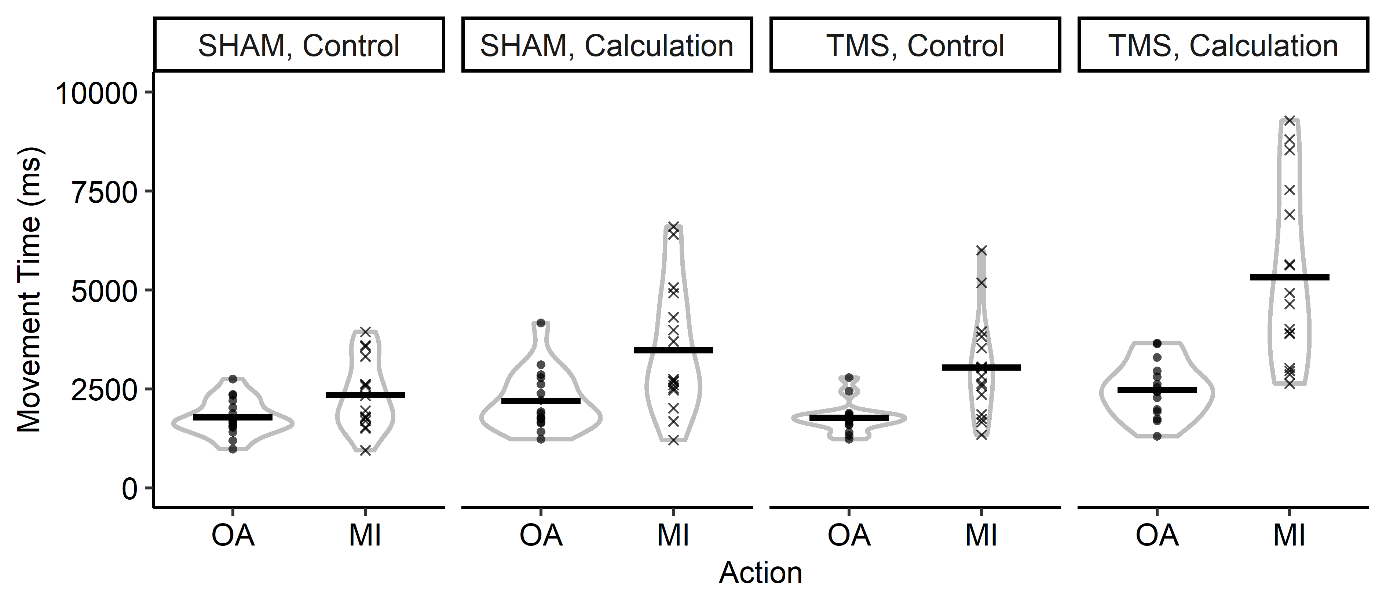

**Figure S4.** Effect of Sham vs. TMS on overt action (OA) and motor imagery (MI) movement times, for the control and the calculation tasks. Horizontal lines represent the group means, individual points represent the means for single participants. Enclosures represent the distribution of the presented data, with wider horizontal sections indicating a larger number of observations.

2.2. Kinematic data

We correlated kinematics and keypresses as in Experiment 1. We excluded 5.7% of the total trials using our outlier criteria. A further 15% of the trials were excluded due to atypical peaks detected by the software. This appears quite high, however, we applied the same cleaning of the data as in Experiment 1 and as stated in our preregistration (<https://osf.io/s82v4/>). Furthermore, this analysis has the purpose of assessing the reliability of the kinematics and the keypresses, and must thus be conducted only on trials for which we are confident that the peak detection was correct, even if we may have been too conservative in our threshold.

As seen in Figure S3, the correlations were high overall, although with more variability between participants than in Experiment 1 and a few participants who had very poor correlations in all three markers of movement time, indicative of a poor coupling between the timing of keypress and the actual movement.

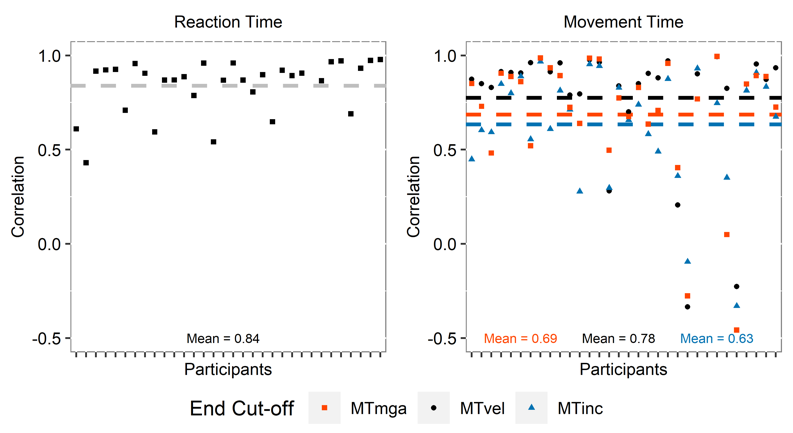

**Figure S5.** Pearson correlation values between the keypress data and the kinematics parameters in Experiment 2. Conventions as in Figure S1.

2.3. Keypress reaction times

As for Experiment 1, we conducted exploratory analysis of the RT data (Figure S4). A model including effects of *TMS, Action,* and their interaction fit the data much better than a null model (λ_adj_ > 1000), indicating that reaction times were longer for the motor imagery condition, and especially so in the TMS group. Adding an effect of *Task* further improved the fit (λ_adj_ > 1000) showing that RT were also higher during the calculation task than in the control condition.

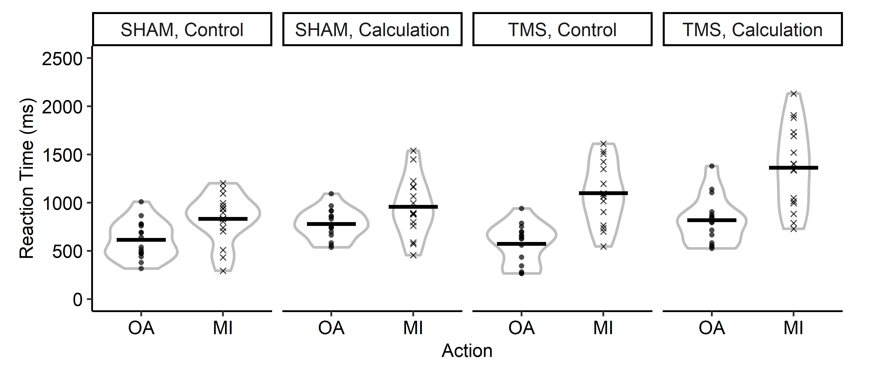

**Figure S6.** Effect of Sham vs. TMS on overt action (OA) and motor imagery (MI) reaction times, for the control and the calculation tasks in Experiment 2. Conventions as in Figure S2.

2.4. Covid Safety Measures

Due to Covid-19 safety precautions, the last 24 participants performed all tasks with a face mask and surgical gloves on (note that one of the excluded participants was part of this “group”). None of the participants reported any additional difficulty in executing the tasks with these safety measures. However, as a control, to examine the effects of Covid safety precautions on keypress movement times, we also conducted an analysis including this variable (*TimelineCovid),* split between “pre-covid” (normal conditions) and “gloves” (post-covid safety restrictions) condition. The ANOVA output thus included a 2 × 2 × 2 × 2 mixed design with *TMS* (TMS/Sham), *Action* (overt action/motor imagery) and *TimelineCovid* (pre-covid/gloves) as between-subjects variables, and *Calc* (calculation/control) as a within-subjects variable.

Data showing the effects of Covid safety measures on keypress movement times in the various conditions are given in Figure S5.

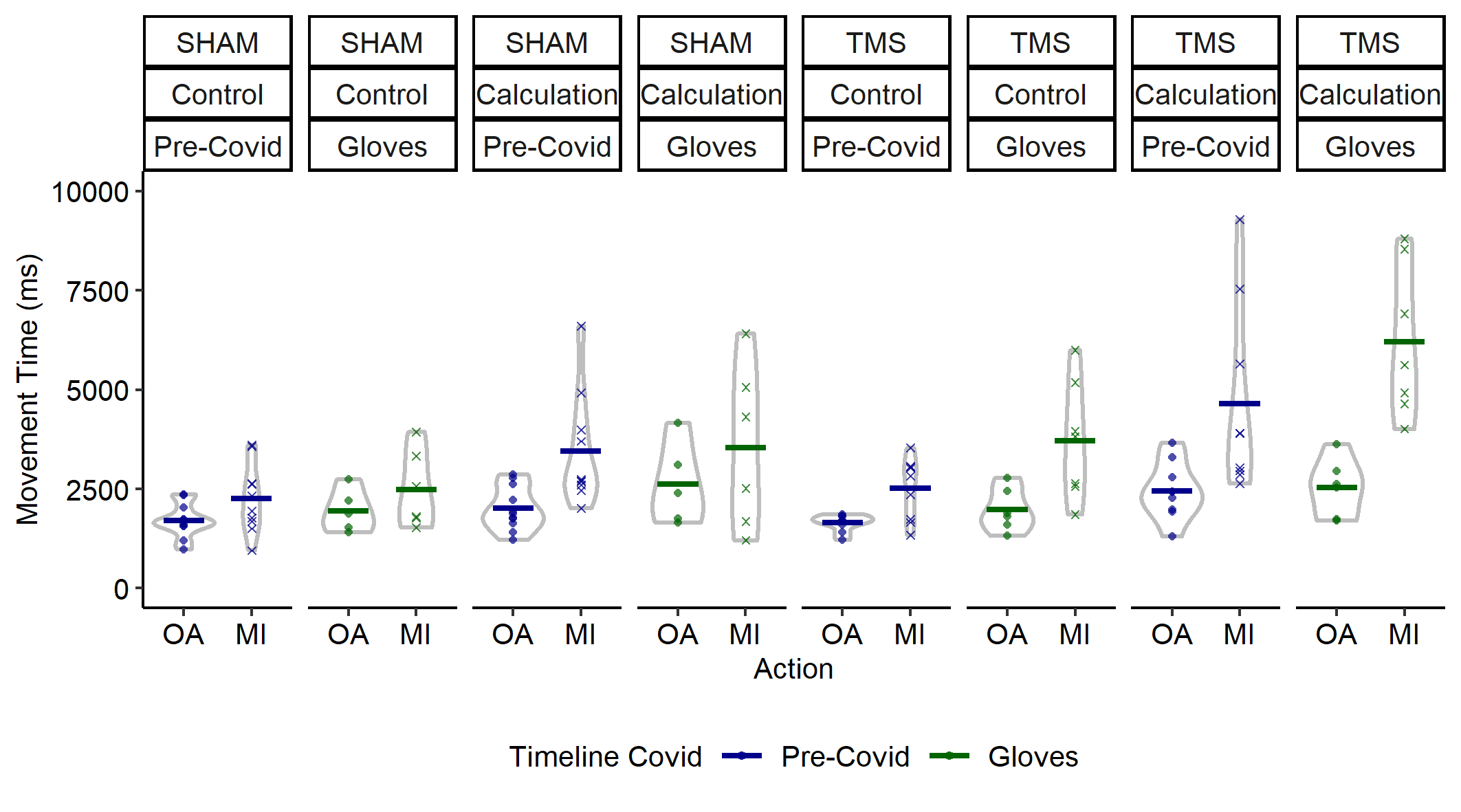

**Figure S7.** Effects of variable TimelineCovid on keypress movement times. Conventions as in Figure S4.

As observed in the ANOVA output below (Table 1), whereas there was some modest evidence that participants following Covid safety measures (n=24) took longer to complete their keypressing than the participants who took part prior to Covid (n=39), λ_adj_ = 1.33, the evidence that the *TimelineCovid* variable did not modulate the effects of the other variables was fairly strong. A null model in which *TimelineCovid* had no interactions with either *TMS* or *Action* (or the three-way interaction) performed better than a model including such interactions (λ_adj_ = 6.88). Similarly, a null model in which *TimelineCovid* had no interactions that included the *Calc* variable performed much better than a model assuming such interactions were present (λ_adj_ = 36.10). Raw data and calculations of likelihood ratios are available on OSF (<https://osf.io/s82v4/>).

**Table 1.** ANOVA output.

| Effect | DFn | DFd | SSn | SSd | F | p |
| --- | --- | --- | --- | --- | --- | --- |
| (Intercept) | 1 | 55 | 959667261 | 110236971 | 478.8 | < .001* |
| TMS | 1 | 55 | 14757144 | 110236971 | 7.36 | .009* |
| ACTION | 1 | 55 | 65310747 | 110236971 | 32.59 | < .001* |
| **TIMELINECOVID** | **1** | **55** | **8689837** | **110236971** | **4.34** | **.042*** |
| CALC | 1 | 55 | 38641800 | 43632689 | 48.71 | < .001* |
| TMS:ACTION | 1 | 55 | 11607071 | 110236971 | 5.79 | .019* |
| **TMS:TIMELINECOVID** | **1** | **55** | **1865084** | **110236971** | **.931** | **.339** |
| **ACTION:TIMELINECOVID** | **1** | **55** | **1540574** | **110236971** | **.769** | **.384** |
| TMS:CALC | 1 | 55 | 3577840 | 43632689 | 4.51 | .038* |
| ACTION:CALC | 1 | 55 | 9338015 | 43632689 | 11.77 | .001* |
| **TIMELINECOVID:CALC** | **1** | **55** | **51152** | **43632689** | **.064** | **.800** |
| **TMS:ACTION:TIMELINECOVID** | **1** | **55** | **3879267** | **110236971** | **1.94** | **.170** |
| TMS:ACTION:CALC | 1 | 55 | 1832519 | 43632689 | 2.31 | .134 |
| **TMS:TIMELINECOVID:CALC** | **1** | **55** | **6644** | **43632689** | **.008** | **.927** |
| **ACTION:TIMELINECOVID:CALC** | **1** | **55** | **6106** | **43632689** | **.008** | **.930** |
| **TMS:ACTION:TIMELINECOVID:CALC** | **1** | **55** | **575210** | **43632689** | **.725** | **.398** |
